# Evolutionary dynamics of multiple games

**DOI:** 10.1101/302265

**Authors:** Vandana R. Venkateswaran, Chaitanya S. Gokhale

## Abstract

Evolutionary game theory has been successful in describing phenomena from bacterial population dynamics to the evolution of social behavior. Interactions between individuals are usually captured by a single game. In reality, however, individuals take part in many interactions. Here, we include multiple games and analyze their individual and combined evolutionary dynamics. A typical assumption is that the evolutionary dynamics of individual behavior can be understood by constructing one big comprehensive interactions structure, a single big game. But if any one of the multiple games has more than two strategies, then the combined dynamics cannot be understood by looking only at individual games. Devising a method to study multiple games – where each game could have an arbitrary number of players and strategies – we provide a concise replicator equation, and analyze its resulting dynamics. Moreover, in the case of finite populations, we formulate and calculate a basic and useful property of stochasticity, fixation probability. Our results reveal that even when interactions become incredibly complex, their properties can be captured by relatively simple concepts of evolutionary game(s) theory.

## Introduction

Evolutionary Game Theory (EGT) [von Neumann and Morgenstern, 1944, Maynard Smith and Price, 1973, Nowak, 2006, Nowak and Sigmund, 2004] has been used to study phenomena ranging from the dynamics of bacterial populations to the evolution of social behavior. In EGT, individuals are cast as players that interact with each other in ‘games’. Games are metaphorical summaries of interactions. For example, in the classical Prisoners’ Dilemma game, individuals can either cooperate or defect, and each pair-wise interaction results in a payoff for the players involved [Nowak, 2006, Nowak and May, 1992]. Over time, players that adopt a certain strategy either perform better than the average population and increase in frequency, or perform worse than the average population and decrease in frequency. Tracking the change in their frequencies over time, EGT can provide insight into the eventual fate of the strategies in a game, e.g. whether they dominate, coexist or go extinct from the population.

However, single games are too simplistic a model. Considerable effort has been done in making them more realistic (with interactions among multiple players and allowing players to adopt strategies from a large set [Ostrom, 1990, 2000]). However, single games fail to satisfactorily capture, for instance, humans interacting in public goods games such as climate change issues [Milinski et al., 2006]. When nations’ leaders discuss strategies to improve the status of global climate, they also need to take into account the interests of the people they are representing. Thus, political leaders are playing at least two games: one with other nations and another within their own nation.

In lions, females defend their territory against invaders by forming a line. Some lionesses always stay at the forefront while others lag behind [Heinsohn and Parker, 1995]. Looking at this territory defense game in isolation, the laggards would be defined as cheaters. Interestingly, the leaders, knowing the identity of the laggards, do not employ any retaliatory strategies (such as Tit for Tat or Pavlov) [Legge, 1995]. The co-existence of the two types would most likely not be evolutionary stable. However, we see stable prides! This puzzle is solved by realizing that territory defense is only one of many games played by the lionesses. In the complete picture, there is division of labor among them, and the laggards could be playing important roles in other games such as maternal care and hunting [Boza and Számadó, 2010, Legge, 1995].

Lastly, a multiple games model in bacterial dynamics can been used to explain the coexistence of avirulent ‘cheaters’ and virulent ‘cooperators’ in populations of the pathogen *S. typhimurium*[Diard et al., 2013]. Likewise, in *Pseudomonas fluorescens* communities, the seemingly destructive cheating cells can promote evolution of collectives [Hammerschmidt et al., 2014]. The dynamics between the microbes constituting the microbiome have been found to be non-linear lending themselves to multiplayer games [Li et al., 2015]. The complete interaction in the holobiont would then be a collection of multiple multiplayer games [?]. In summary, real world interactions cannot be described by single games [Tarnita et al., 2009] and as mentioned earlier, one particular use of multiple games is tackling the evolution of division of labor [Rueffler et al., 2012, Gazda et al., 2005, Wahl, 2002, Kerr et al., 2002].

Previous studies on multi-game dynamics (MGD) (Fig. 1) have shown that a combination of games with more than two strategies cannot be separated into it’s constituent single game dynamics (SGD) [Hashimoto, 2006]. However, these results are restricted to two-player games. When more players are involved, different dynamics emerge [Pacheco et al., 2009, Gokhale and Traulsen, 2010, Peña, 2012] A complete picture of MGD, where multiple players are involved, is lacking. If multiple players are involved, then can the MGD be decomposed back into its constituent SGDs? If yes – the conclusions drawn from individual games are valid. If not – it will be necessary to use MGD to obtain realistic results.

**Figure 1:**
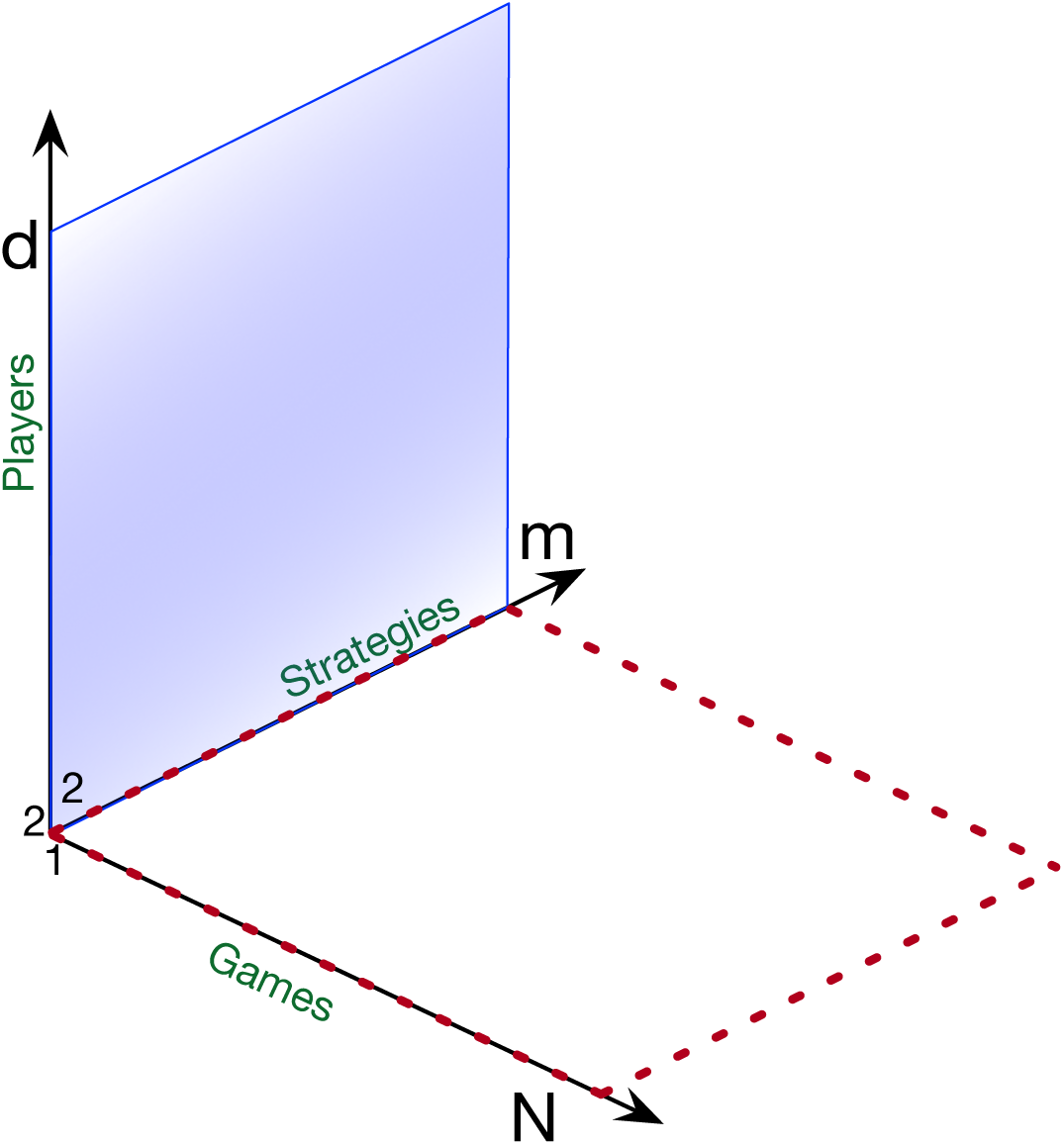
Scope of this study. Typical evolutionary game dynamics focuses on two player games with two strategies. Extensions to multiplayer games (*d*) and multiple strategies (*m*, solid blue rectangle) expands the domain of study to public goods games and other social dilemmas. However this is still limited to a single game. Hashimoto [2006] has extended two player-multi-strategy games in a novel direction of multiple games (*N*, dotted red rectangle). Our work generalizes this approach and develops a method for analyzing multiple games, where each involved game could be a multiplayer (and multi-strategy) game. Thus, this approach enables us to study the entire space of multiple games (*N*) with multiple strategies (*m*) consisting of multiple players (*d*).

To answer this question, we enhance the MGD to look at combinations of multiplayer games, and provide an analytical framework for analyzing an ensemble of games in a tractable manner. We present a complete and general method to study multiple games with many strategies and players, all at once (Fig. 1). When the games have more than two strategies, we find that they cannot be separated back to their SGDs, in line with previous findings. Interestingly, however, we find a dependency on the initial conditions (i.e., the initial frequencies of each strategy). For certain initial conditions, one may still be able to capture the SGDs from their MGD.

## Model and Results

### Single game dynamics (SGD)

Two player games with two strategies have been studied extensively, both in infinite as well as finite populations. A game between two individuals can be represented by the following interaction matrix,

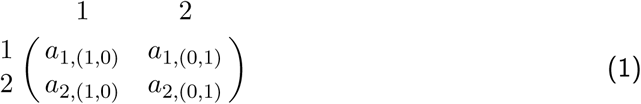

The two individuals are represented by a row and a column respectively and each can adopt one of the two strategies 1 or 2. We write the elements of the matrix in the form *a*_*i*,*α*_, where *i* is the strategy of the focal player. The vector *α* is written as *α* = (*α*_1_, *α*_2_) where *α*_1_ (number of individuals of strategy 1 in the column) and *α*_2_ (number of individuals of strategy 2 in the column), together represent the group composition. In a 3-player game with two strategies, a payoff matrix entry, say *a*_2,(1,1)_, where *α*_1_ = 1 and *α*_2_ = 1, will correspond to a focal player with strategy 2 interacting with two other players with strategies 1 and 2, respectively.

The average payoff obtained from the game is the reproductive success of that strategy [Maynard Smith, 1982]. This analysis has been extended to interactions having *multiple* strategies [Wu et al., 2011] as well as *multiple* players [Broom et al., 1997, Broom, 2003]. To make our notation clear, we illustrate a payoff matrix for a multiplayer (*d* player) game with two strategies as,

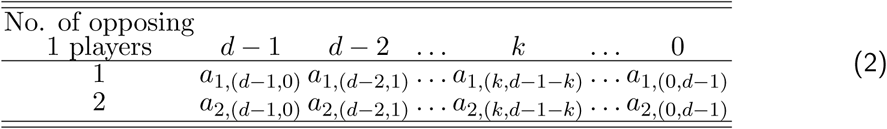

Even while extending the number of strategies, the dynamics of this complicated system can still be analyzed by the replicator dynamics [Hofbauer and Sigmund, 1998, Schuster and Sigmund, 1983]. For a *d* player game with m strategies, the replicator dynamics is given by a set of m differential equations

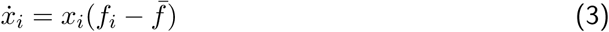

where *x_i_* is the frequency of strategy *i*, and *f_i_* is the fitness of the strategy *i* (see Supplementary Information (SI) text). The average fitness of the population is given by
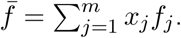 This simple evolutionary game framework has been used to describe a wide range of phenomena from chemical reactions of prebiotic elements to the evolution of social systems [Komarova, 2004].

While this extension to multiple players and strategies is not trivially obtained [Gokhale and Traulsen, 2011], it still belongs to the domain of a single game. To make EGT models more realistic, interactions which have differential impacts on fitness need to be taken into account. Therefore, we now incorporate multiple games and measure their cumulative impact on individual fitness.

### Multi-game dynamics (MGD)

Individuals may employ different strategies in various games (e.g., division of labor scenarios [Wahl, 2002]) and their (average) payoffs will depend on their performance on all such games. Switching between such socially driven games is realistic and not only a matter of theoretical interest but has been experimentally explored as well [Wedekind and Milinski, 1996].

To contrast multi-game dynamics (MGD) with the previously discussed single game dynamics (SGD), consider a simple example of two, 2 × 2 games:

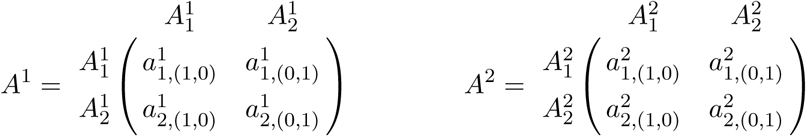

Combining the strategies from the above two games results in four categories of individuals (Fig. 2). The frequencies of the four categories are given by *x*_11_, *x*_12_, *x*_21_ and *x*_22_ where the first and second positions (in the subscript) denote the strategies adopted in games 1 and 2, respectively (Fig. 2). For a combination of *N* games, in principle, each game *j* can be described by a payoff matrix *A^j^*. Each game *j* could be a *d_j_* player game with *m_j_* number of strategies. The categorical frequencies would then be given by *x*_*i*_1_*i*_2_…*i_j_*…*i_N_*_ where *i_j_* is the strategy being played in game *j*.

**Figure 2:**
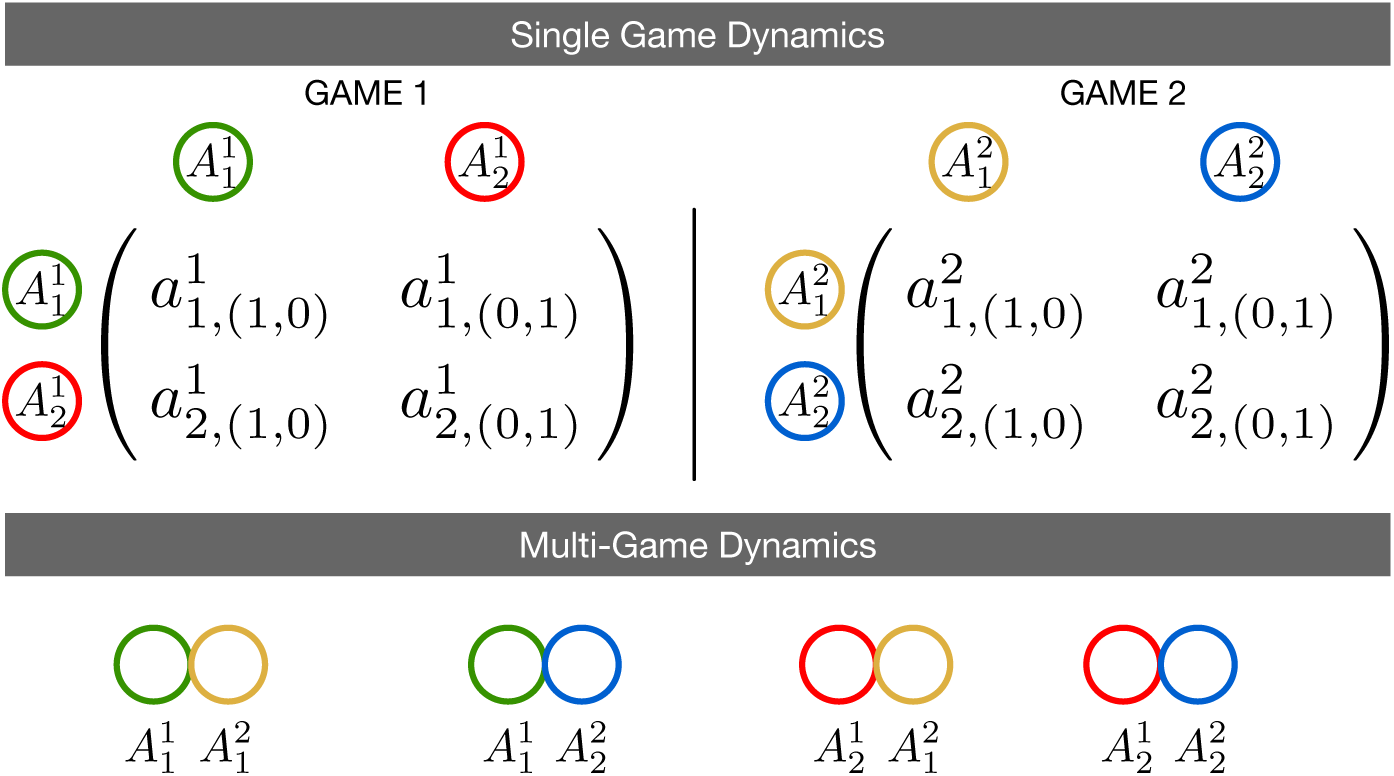
From Single Game Dynamics to Multi-Game Dynamics. The population after combination is divided into four types: playing strategy 1 in game *A*^1^ and game *A*^2^, strategy 1 in *A*^1^ and 2 in *A*^2^, strategy 2 in *A*^1^ and 1 in *A*^2^. And finally, strategy 2 in *A*^1^ and *A*^2^. Thus,we have four types of strategies,
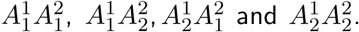 Their respective frequencies are *x*_11_, *x*_12_, *x*_21_ and *x*_22_. Since there are four ‘categorical types’, we can project the dynamics on an *S*_4_ simplex.

The frequencies of the individual strategies for all *N* games can be written down as,

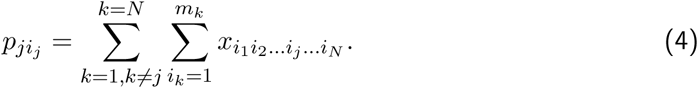

Using this individual strategy frequency for a game *j*, the fitness of strategy *i_j_* is given by,

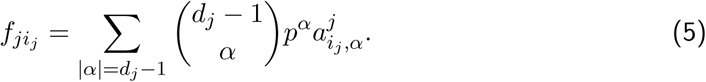

As before, *α_m_j__* is the number of strategy *m_j_* players. Using multi-index notation, we have *α* = (*α*_1_, *α*_2_, …, *α_m_j__*) which gives us the multinomial coefficient, with the absolute value |*α*| = *α*_1_ + *α*_2_ + … + *α_m_j__* and the power
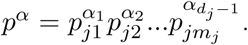 The average fitness of the population is given by, *ϕ_j_* = (**pf**)_*j*_ (see SI text). Putting all this information together, we can write down the time evolution of all the categorical strategies as,

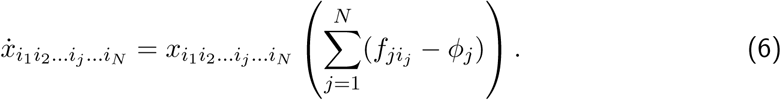

This system of equations is reminiscent of the replicator equation for the SGD. The summation in the MGD replicator equations is due to an assumption of additive fitness effects from all games [Hashimoto, 2006]. In the following sections we will explore the use of this formulation for multiple games where each game can have a different number of players. Through the examples of specific cases, we aim to highlight the general principles of multiple games.

### Two player game(s) with multiple strategies

In case of two player games with two strategies, Cressman et al. [2000] showed that the SGD can be separated from the MGD. The dynamics lies on the generalized invariant *Wright manifold* [Hofbauer and Sigmund, 1998] in the *S*_4_ simplex which is given by *W_K_* = {*x* ∈ Δ^4^ | *x*_11_*x*_22_ = *Kx*_12_*x*_21_} for *K* > 0. All the trajectories in the simplex depicting the MGD fall onto an attractor given by a line (ES set) on *W_K_*. However, previous results [Hashimoto, 2006] show that for more than two strategies, the MGD cannot be separated even into a linear combination of the constituent SGDs unless they are on *W_K_*. We are clearly looking at higher dimensions and the space is dense with various manifolds. It is important to know on which manifold the initial conditions are, for only if they start from *W_K_*, will the system state end on *W_K_*.

### Multiplayer game(s) with multiple strategies

In combinations containing multiplayer games, frequency feedback between strategies is possible. Moreover, as discussed in the beginning, an individual can take part in different interactions. A lioness can be part of forming the defensive line (tragedy of the commons) and hunting (stag-hunt game). Strategies in game 1 would be *Cooperator*, *Defector*, *Loner* etc. Strategies in game 2 could be *Wing*, *Centre* and so on. Thus
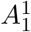 need not be the same as
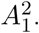 Using our framework, we can model the combined dynamics of several games that an individual plays where each game can have completely different strategy sets.

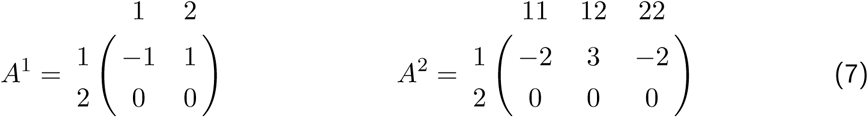

To illustrate games with two strategies, we shall use the payoff matrices shown in (7). Here, *A*^1^ is a two player coexistence game and *A*^2^ is a three player game. In Game *A*^2^, the values *a*_1,(*k*,*d*–1–*k*)_ – *a*_2,(*k*,*d*–1–*k*)_ and *a*_1,(*k*+1,*d*–*k*)_ – *a*_2,(*k*+1,*d*–*k*)_ have different signs for all *k*. Thus, we have two interior fixed point solutions: a stable and an unstable. The exact solutions for the two SGD’s in (7) are
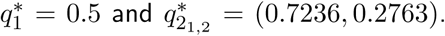 The result of combining these games i.e. their MGD, is shown in Fig. 3.

**Figure 3:**
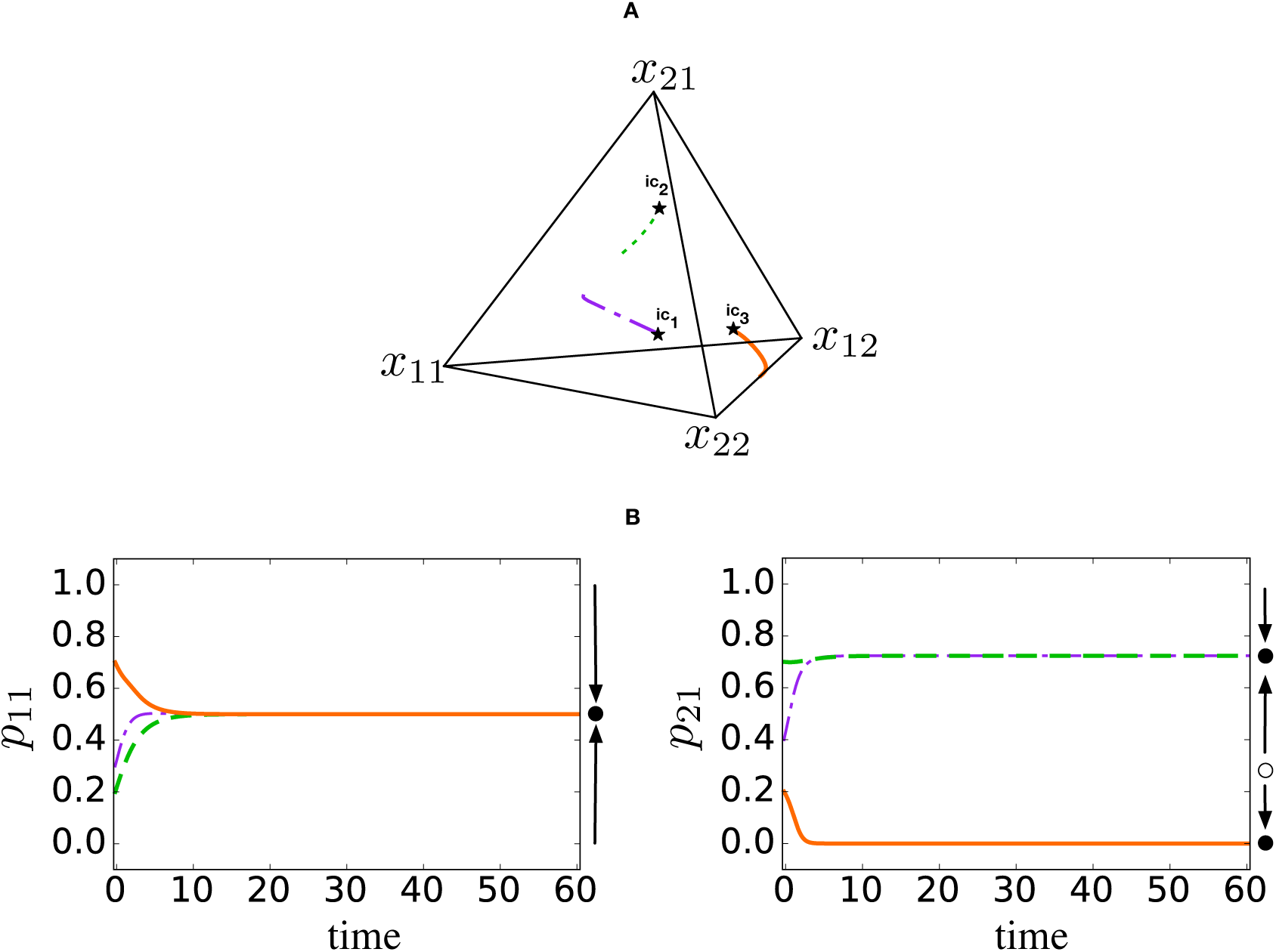
(A) This *S*_4_ simplex contains the Multi-Game Dynamics of the combination of the 3-player and 2-player games in (7). The vertices are made up of the four ‘categorical strategies’. The asterisks depict the initial conditions (*ic*_1_, *ic*_2_ and *ic*_3_) of the three trajectories thats are plotted here. (B) In the multi-game dynamics, *p*_11_ (playing strategy 1 in game 1) converges to **q_1_** = 0.5 which is the equilibrium solution for strategy 1 in game 1. If we start above the unstable equilibrium solution for game 2 i.e **q_2_2__** = 0.2763932, then *p*_21_ (playing strategy 1 in game 2) converges to **q_2_l__** = 0.7236068 which is the stable equilibrium solution for game 2. For trajectories that commence below the unstable equilibrium, strategy 1 goes to extinction. Clearly, *p*_12_ = 1 – *p*_11_ and *p*_22_ = 1 – *p*_21_. The initials conditions for {*x*_11_, *x*_12_, *x*_21_ and *x*_22_} used in these plots are : *ic*_1_ = {0.2, 0.1, 0.2, 0.5}, *ic*_2_ = {0.1, 0.1, 0.6, 0.2} and *ic*_3_ = {0.1, 0.6, 0.1, 0.2}.

Next, we shall look at an example where one game, say, *A*^1^, has three strategies. Let *A*^1^ to be a Rock-Paper-Scissor type game as shown in the first payoff matrix in 8.

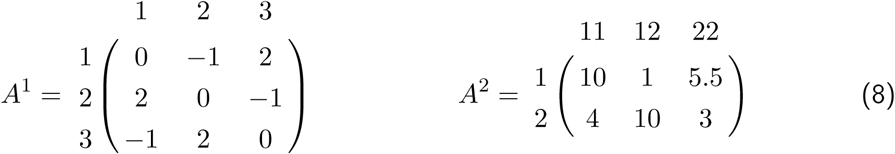

Since the determinant of the matrix is positive, the trajectories starting from any initial condition will converge to a unique stable equilibrium. The other game, *A*^2^, as shown in (8) is a three player game similar to the one used before in (7). In the SGDs of these games, the interior solution for Game *A*^1^ is **q_1_** = (1/3,1/3,1/3). For Game *A*^2^, the equilibrium solutions are **q_2_l__** = 0.127 (stable) and **q_2_2__** = 0.740 (unstable). The outcomes of their MGD will be on an *S*_6_ simplex. Since it is not possible to show this simplex, the importance of using *p_ji_j__*. is clear as we use these now to compare the MGD with their SGDs. The projections are as shown in Fig. 4.

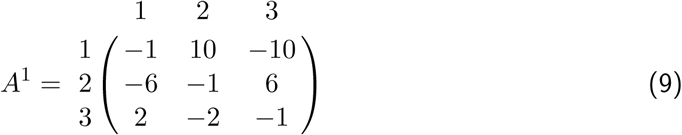

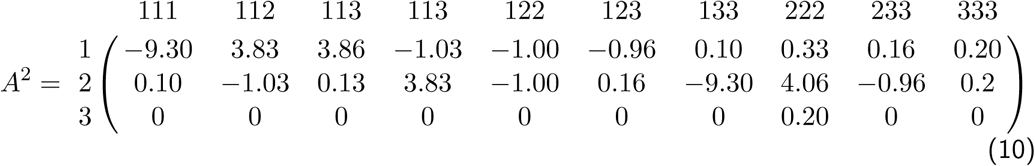

**Figure 4:**
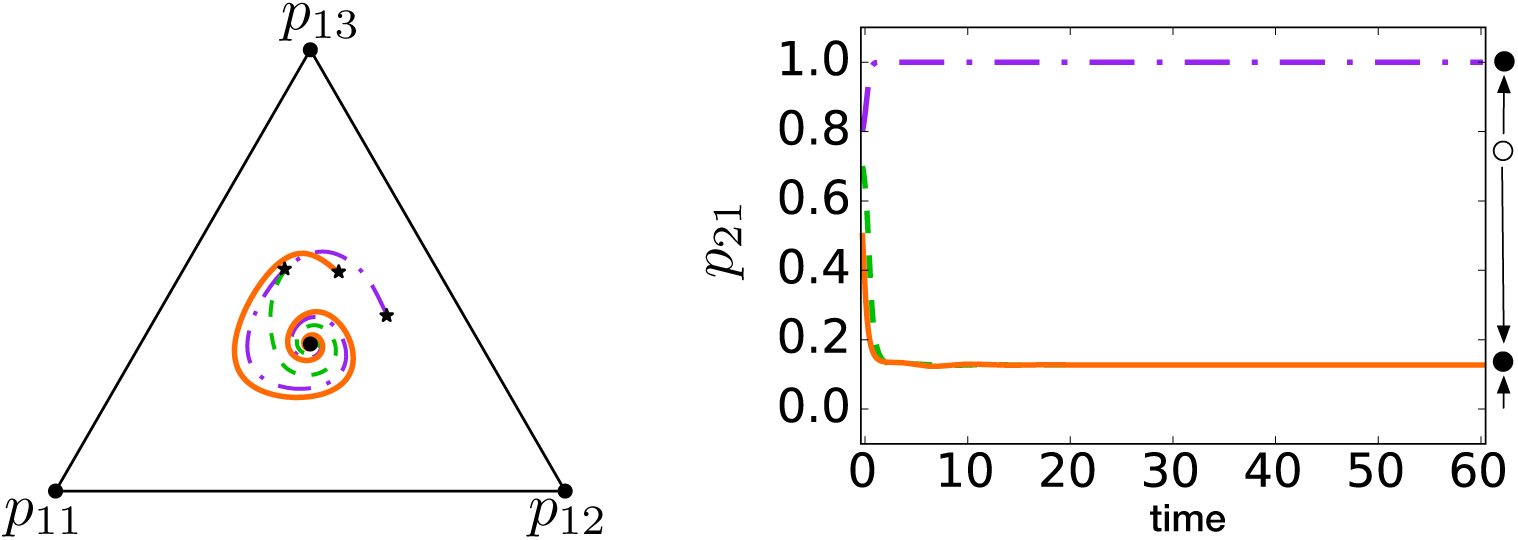
When there are three strategies in one game and two strategies in the other, six “categorical types” are possible in their multi-game dynamics. The MGD will be on an *S*_6_ simplex. Avoiding a five dimensional figure, we retrieve the distribution of frequencies of strategies in the SGDs from the MGD which is what we require to compare the SGDs and MGDs. The asterisks in the triangular *S*_3_ simplex denote the positions from where the trajectories begin (initial conditions). Retrieving the distribution of frequencies of strategies in game *A*^1^, all trajectories converge to the equilibrium solution **q_1_** = (1/3,1/3,1/3) and in game *A*^2^, the trajectories that begin from below the unstable equilibrium **q_2_2__** = 0.740 converge to the stable equilibrium solution **q_2_l__** = 0.127. The initials conditions used for {*x*_11_, *x*_12_, *x*_21_, *x*_22_, *x_31_* and *x*_32_} are : *ic*_1_ = {0.3, 0.1, 0.1, 0.05, 0.4, 0.05}, *ic*_2_ = {0.4, 0.1, 0.2, 0.1, 0.1, 0.1} and *ic*_3_ = {0.2, 0.3, 0.1, 0.1, 0.2, 0.1}.

Finally, we shall illustrate a case of having three strategies in both games (shown in matrices 9 and 10). *A*^1^ is a Rock-Paper-Scissor game like the one discussed in the previous example. *A*^2^ is a 4-player three strategy game used previously in Gokhale and Traulsen [2010]. In the SGDs of the individual games, *A*^1^ has a stable equilibrium solution **q_1_** = (1/3,1/3,1/3) and since *A*^2^ is a 4-player three strategy game, it has (*d* – 1)^(*n*–1)^ = 3^2^ = 9 interior equilibrium solutions : four stable, one unstable and four saddle points. The resulting Multi-Game Dynamics is shown in Fig. 5.

**Figure 5:**
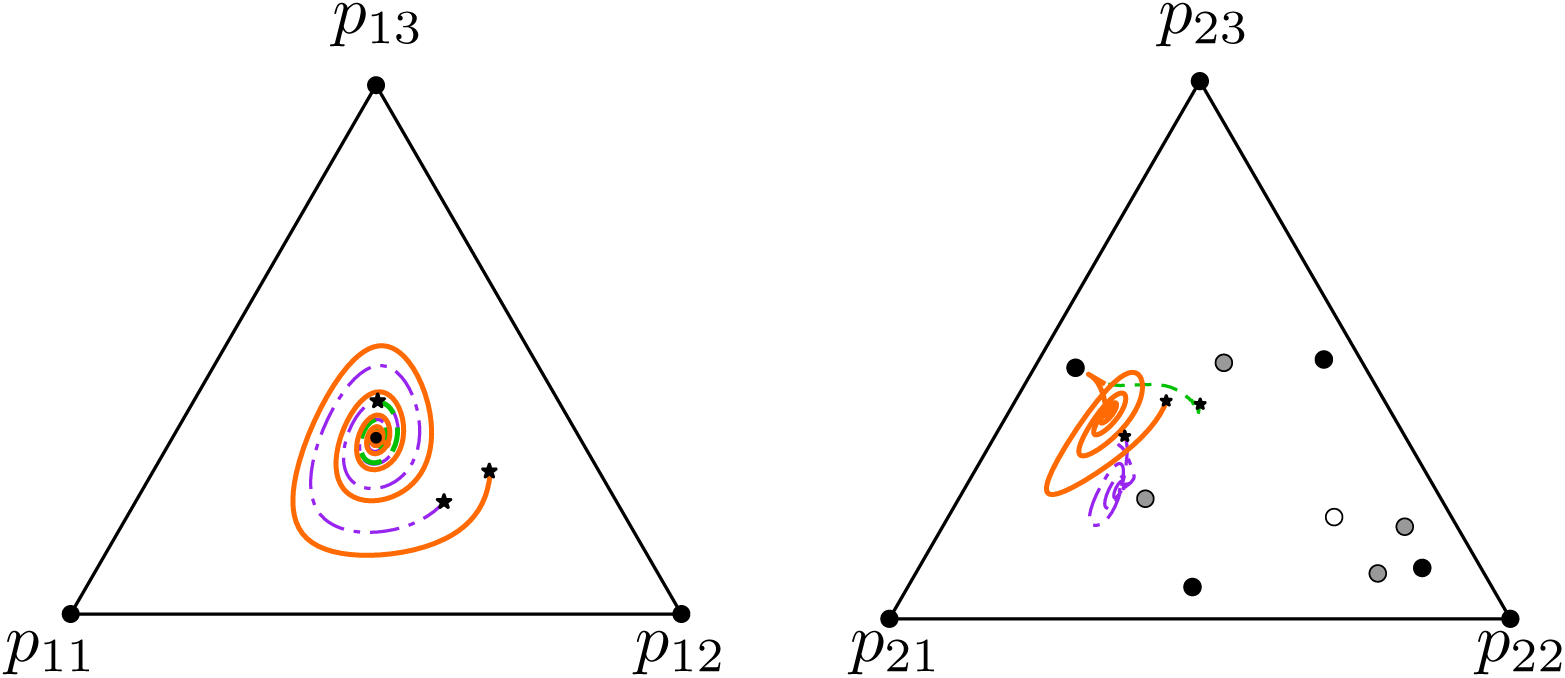
When both games contain three strategies nine categorical types are possible. The MGD would be in an *S*_9_ simplex. As discussed in Fig. 4, since we avoid an eight dimensional figure, we retrieve the distribution of frequencies of strategies in the SGDs from the MGD which is what we require to compare the SGDs and MGDs. The asterisks in the triangular *S*_3_ simplex denote the initial conditions. The triangular markers are the final position of the trajectories. The black, grey and white solid circles are the stable, saddle and unstable interior equilibrium solutions in the SGDs. While retrieving the distribution of frequencies of strategies in the SGDs from the MGD, we see that not all trajectories converge to the equilibrium solutions of the SGDs. When both games have more than two strategies, initial conditions matter. For few initial conditions, we can decompose the multi-game into its constituent games and for others, we cannot. The initials conditions used for *x*_11_, *x*_12_,*x*_13_, *x*_21_, *x*_22_, *x*_23_,*x*_31_, *x*_32_ and *x*_33_ are : *ic*_1_ = {0.01, 0.166, 0.038, 0.002, 0.176, 0.102, 0.3251, 0.111, 0.070}, *ic*_2_ = {0.2, 0.1, 0.1, 0.1, 0.1, 0.1, 0.1, 0.1, 0.1} and *ic*_3_ = {0.176, 0.066, 0.024, 0.002, 0.176, 0.002, 0.225, 0.111, 0.218}.

For multiplayer games, we perform a similar study as in two player games. The MGDs can be separated into SGDs if both the games have only two strategies (Fig. 3). The expression for *W_K_* though, would have higher order terms. Thus, the attractor may no longer be a line, but instead a curve *W_K_* in a higher dimensional space. We performed an analysis where only one game has two strategies (Fig. 4) and here too the MGDs can be separated into their integral SGDs. However, while considering more than two strategies in both games (Fig. 5), the MGDs cannot always be trivially separated into their constituent SGDs. As in Hashimoto [2006], it becomes important to look at the initial conditions. Some trajectories converge to the fixed point solutions of the SGDs, while many others do not. Table 1 provided in the appendix contains a condensed description of the effect of initial conditions.

### Finite population

Evolutionary dynamics in finite populations has the potential of having qualitatively different dynamics than their deterministic analogues [Nowak et al., 2004]. In finite populations the size of the population controls the balance between selection and drift with small populations showing higher levels of stochasticity.

We use a birth-death Moran process to model a finite population of size *Z* in our framework [Nowak et al., 2004, Traulsen and Hauert, 2009]. An individual is chosen (proportional to its fitness) to reproduce an identical offspring. Another individual is chosen randomly for death. Thus the total population size remains constant. Fitness, as measured before, is a function of the average payoffs. Besides the population size, we can control the effect of the game on the fitness via a payoff to fitness mapping. The mapping could be a linear function *f* = 1 – *w* + *wπ* where *w* is the selection intensity [Nowak, 2006]. If *w* = 0, selection is neutral whereas for *w* = 1 selection is strong and the payoff determines the fitness completely. However, since negative fitnesses in this framework are meaningless, there are restrictions on the range of *w*. Alternatively, to avoid this restriction, we can map the payoffs to fitness using an exponential function *f* = *e^wπ^* [Traulsen et al., 2008]. The fixation probability of strategy 1 in a SGD for a d-player game, under weak selection, is given by [Gokhale and Traulsen, 2010],

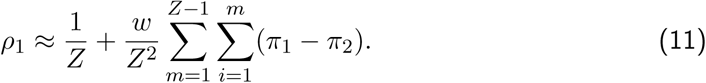

where *π_i_* is the average payoff of strategy *i*. We have extended this to multiple games. In order to do this, we define what we mean by fixation probability in multiple games. The strategies in a multiple game are the categorical ones. For instance, a two game system with each game containing two strategies, has four categorical strategies as shown in Fig. 2. In a population of size *Z* playing *N* multi-strategy *d*-player games, if *γ* is the number of individuals of the categorical type
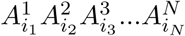 on an edge, then the number of individuals of type
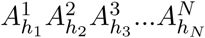 on the other vertex of that edge will be *Z* – *γ*. The average payoff is given by (see SI text),

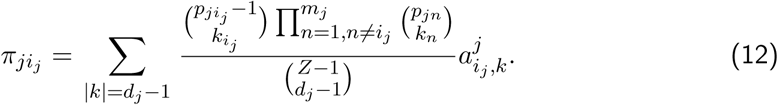

The above expression is utilized to calculate the fitnesses of the categorical types *F*_*i*_1_*i*_2_*i*_3_…*i_N_*_ and *F*_*h*_1_*h*_2_*h*_3_…*h_N_*_ (see SI text) and using these expressions, the fixation probability of type
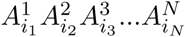 fixating in population of
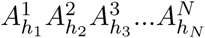 becomes equal to,

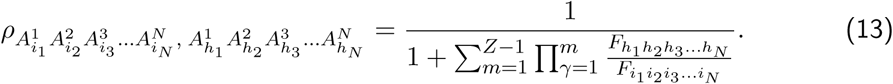

We make pairwise comparisons between all categorical types (all the edges of the *S*_4_ simplex in Fig. 2 containing the MGD). Using these comparative fixation probabilities we can determine the flow of the dynamics over pure strategies as shown in Fig. 6 (see SI text).

**Figure 6:**
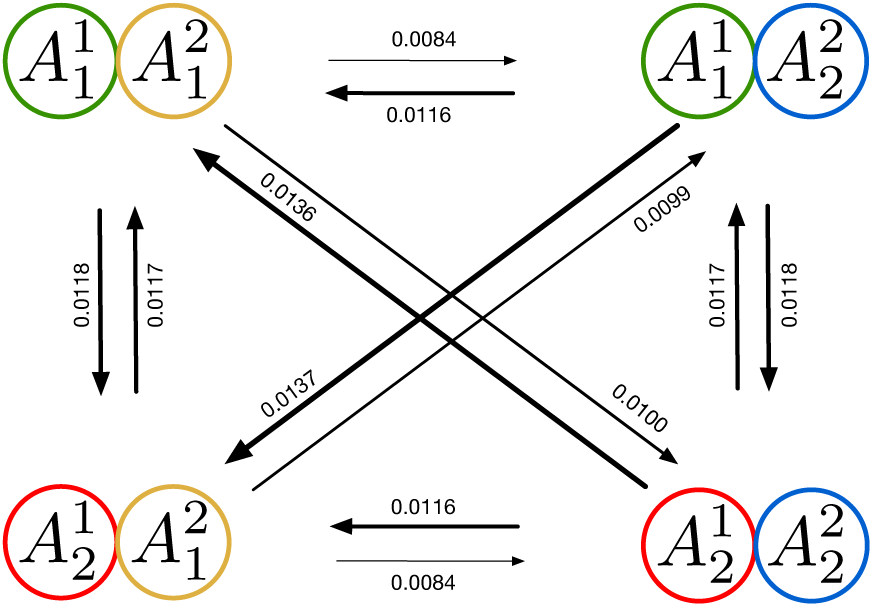
Figure showing the direction of selection and strength according to fixation probabilities between the vertices in a tetrahedron (which contains the MGD of games shown in matrices 7) shown in Fig. 3. The width of the arrows correlate with the magnitudes of the fixation probabilities. Here selection intensity *w* = 0.01 and population size *Z* = 100. It has been assumed that both the games have the same selection intensity and hence the average payoffs have been added first and then the mapping has been performed. For different mappings for the two games, see SI. (Result I). For the edges where one of the games does not change (e.g.
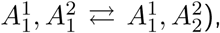 only one of the game (here game 2) matters and hence the fixation probabilities are the same as if *only* one game.

Instead of merely looking at the fixating probabilities of certain types or strategies in a game, we have expanded the method for analyzing the ‘categorical types’ in the multi-game dynamics (see SI text). Therefore, one can determine the dynamics of entities playing a combination of different roles (strategies) in various interactions (games).

## Conclusion

Nature is composed on many interactions (games). The games consist of different players and strategies. And one player in its time plays many parts (in various games).

We devised a method to combine the various multiplayer multi-strategy games that individuals play at a certain period to incorporate the observed complexity while modelling games that biological entities play; to advance further into creating more realistic models. Taking more than two strategies into account represents situations such as the three strategy rock-paper-scissor like games that *E.coli* play in addition to a public goods game [Wakano et al., 2009, Kerr et al., 2002].

While biological and social analogies of multiplayer evolutionary games can be found aplenty, the case for considering multiple games is strong. The gut microbiota is a complex system which is capable of showing a variety of stable states, often a dynamic stability [Li et al., 2015, Abedon, 2008]. The different microbes within the gut community definitely interact in a variety of ways within themselves but each also interacts with the host in a unique manner. Within species and between species interactions, together, have the potential to dictate the evolutionary course of all involved species [?]. These interactions can certainly be interpreted as multiple games, each with a number of strategies and (immensely) multiplayer games. On the population genetics level, as an extension to the work by Traulsen and Reed [2012], multiple games and multi-strategies can be seen as multiple loci with several alleles. The case for two loci and two strategy games has been investigated by Cressman et al. [2000] while the three strategy games by Hashimoto [2006]. Since we consider more than two players, our work can be extended to investigate polyploidy as well [Han et al., 2012].

In a nutshell, from the analyses that we performed, the outcomes from multiplayer two strategy games are similar to Cressman et al. [2000]’s results where the MGD can be characterized by the separate analysis of the individual games. However, when the games have at least three pure strategies, different dynamics emerge. For such cases, a fully comprehensive study of the initial conditions is a potential future work. It would be interesting to see what fraction of them end up converging to the equilibrium solutions of the individual games. Even though complicated dynamics can still be captured by the relatively simple replicator like equations, vast domains in the multiple games space remain unexplored.

## APPENDIX

**Table 1:** When all games involved consist of more than two strategies, initial conditions of the trajectories matter. While retrieving the SGDs from the MGDs, only a fraction of the trajectories converge to the fixed point solutions of the individual SGDs i.e. *p*_11_ = *q*_1_ and *p*_21_ = *q*_2_ only for certain initial conditions. In other words, while extracting the SGDs from the MGDs, they do not behave like the individual games for all initial conditions and therefore, we cannot decompose the MGD into its inherent games. To analyze the sensitivity of initial conditions, they were rounded up to 10 decimal places, first. Later, we allowed up to 100 decimal places. The two player games used in this table are from Cressman et al. [2000] and Hashimoto [2006] and the multiplayer games are the ones discussed in the main article of this paper. In all examples involving R-P-S games, the check for convergence to the internal equilibrium 0.33 was done by rounding it up to four decimal places i.e. 0.3333.

| Games | Initial conditions rounded up to 10 decimal places | Initial conditions rounded up to 100 decimal places |
| --- | --- | --- |
| $(2 \times 2) + (2 \times 2)$ | All trajectories converged | All trajectories converged |
| $(3 \times 3) + (3 \times 3)$ | All trajectories converged | 0.46% of total trajectories converged |
| $(2 \times 2) + (2 \times 2 \times 2)$ | All trajectories converged | All trajectories converged |
| $(3 \times 3) + (2 \times 2 \times 2)$ | All trajectories converged | All trajectories converged |
| $(3 \times 3) + (3 \times 3 \times 3 \times 3)$ | No trajectory converged | No trajectory converged |

## SUPPORTING INFORMATION

### 1 Infinite population

#### 1.1 Single Game Dynamics (SGD)

##### A two player replicator approach

Consider a 2 × 2 (two player two strategy) payoff matrix (14) : There are two players and each of them can adopt two strategies The two types of strategies they could employ are 1 and 2 and their respective frequencies are *x*_1_ and *x*_2_.

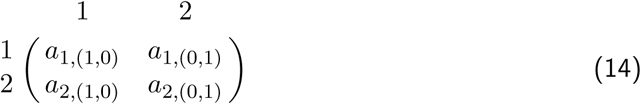

In matrix 14, we write the elements in the form *a*_*i*,*α*_, where *i* is the strategy of the focal player. *α* (using multiindices notation) is a vector written as *α* = (*α*_1_, *α*_2_). *α*_1_ and *α*_2_ together represent the group composition. The average payoffs of the two strategies are given by *f*_1_ = *a*_1,(1,0)_*x*_1_ + *a*_1,(0,1)_*x*_2_ and *f*_2_ = *a*_2,(1,0)_*x*_1_ + *a*_2,(0,1)_*x*_2_. The replicator equation Eq. (15) [Hofbauer and Sigmund, 1998, Nowak, 2006] describes the change in frequency *x_i_* of strategy *i* over time.

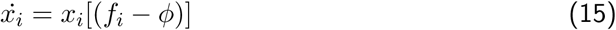

where *f_i_* is the fitness of strategy *i* and *ϕ* is the average fitness. For an infinitely large population size we have *x*_1_ = *x*, *x*_2_ = 1 – *x* Thus the replicator equation for the change in the frequency of stratey 1 is,

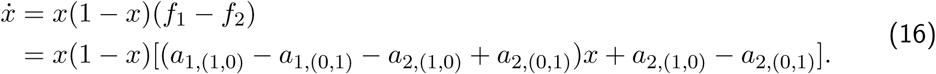

Apart from the trivial fixed points (*x* = 0 and *x* = 1), there is an internal equilibrium given by,

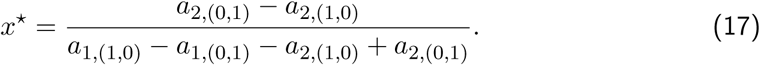

##### Multiplayer games

We now extend the dynamics to multiplayer games [Gokhale and Traulsen, 2014]. The payoff matrix (18), represents a three player (*d* = 3) two strategy (*n* = 2) game; a 2 × 2 × 2 game.

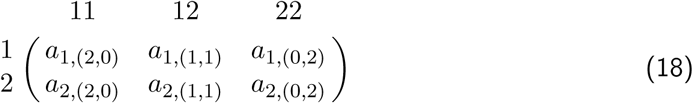

The rows correspond to the focal player. Focal player interacting with two other players, both with strategy 1 will receive a payoff *a*_1,(2,0)_. While interacting with one strategy 1 player and another strategy 2 player, he will get *a*_1,(1,1)_. Interacting with two other strategy 2 individuals, the payoff is equal to *a*_1,(0,2)_. Assuming that the order of players does not matter, the average payoffs (or in this case, the fitnesses) will be,

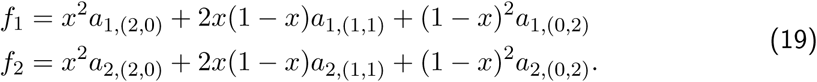

The replicator equation in this case is given by,

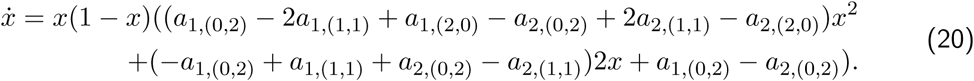

The quadratic *x*^2^ term in Eq. (20) can give rise to a maximum of two interior fixed points. In general, for a *d*-player two strategy game, the replicator equation can result in *d* – 1 interior fixed points (maximum). For an n strategy *d*-player game, the maximum number of internal equilibria is (*d* – 1)^(*n*–1)^ as shown in Gokhale and Traulsen [2010].

#### 1.2 Multi Game Dynamics (MGD)

##### Linear combination of two 2 × 2 games

To start looking into the dynamics of combinations of games i.e. Multi Game Dynamics (MGD) in contrast with the Single Game Dynamics (SGD), consider the example: two games with two strategies in each. Let the payoff matrix of Game 1 and Game 2 be,

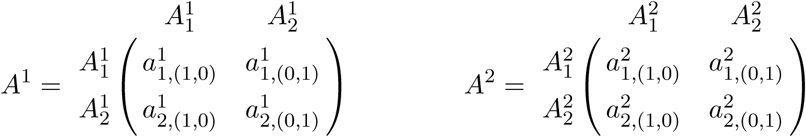

The individuals can be partitioned into four classes. Individuals playing strategy 1 in game *A*^1^ and game *A*^2^, strategy 1 in *A*^1^ and 2 in *A*^2^, strategy 2 in *A*^1^ and 1 in *A*^2^, and strategy 2 in *A^1^* and *A*^2^. So, there are four types of strategies,
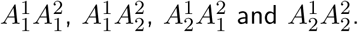 We refer to them as “categorical types”. Their respective frequencies are written as *x*_11_, *x*_12_, *x*_21_ and *x*_22_. We shall now use a new notation, *p_i_j_j_* or playing strategy *i_j_* in game *j*, which is just a variable transformation that can be written as (here, *i_j_* ∈ {1,2} and *j* ∈ {1,2}),

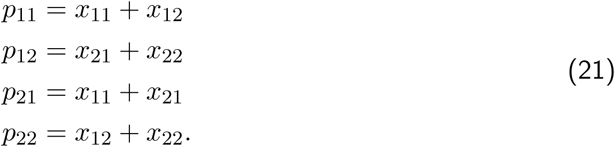

The fitnesses for playing strategy *i_j_* in game *j* can be written out as,

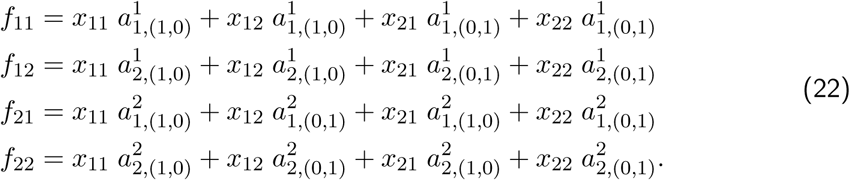

A crucial assumption here is that the effective average payoff is a linear composite of the constituent games. The replicator dynamics will be given by the following set of coupled different differential equations

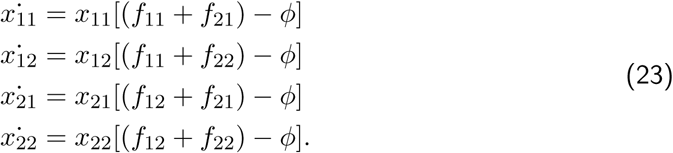

The average fitness *ϕ* is given by,

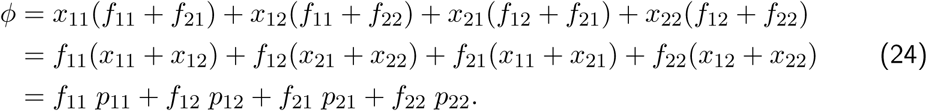

### 2 Finite population

#### 2.1 Single game dynamics

In a population of size Z consisting of strategy 1 and 2 players, the probability that one of the strategies, say 1, fixes, is given by the fixation probability *ρ*_l_. An individual is chosen proportional to its fitness to reproduce an identical offspring. Another individual is chosen randomly and discarded from the group. Therefore, the group size is kept at a constant value *Z*. Fitness of a strategy *s* can be a linear function of its average payoff *π_s_* i.e *f_s_* = 1–*w*+*wπ_s_*. In a population that has *i* strategy 1 players, the fitnesses can be used to calculate the transition probabilities
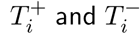 for the number of type 1 players to increase and decrease by one, respectively.

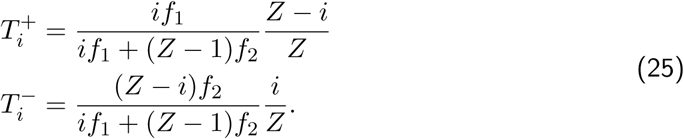

With probability
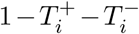 the system does not change. Using the transition probabilities, the fixation probability can be calculated [Nowak, 2006, Traulsen and Hauert, 2009] to be,

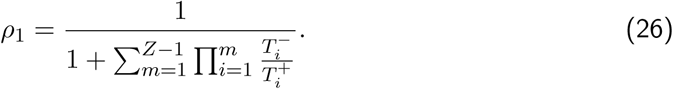

Since
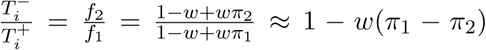 for selection intensity *w* ≪ 1 i.e. weak selection. Therefore,

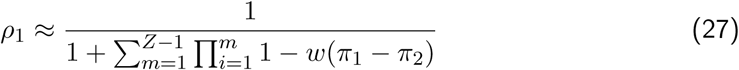

For a *d*-player game, the payoffs are obtained using a hypergeometric distribution given by,

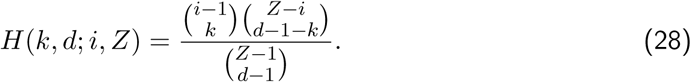

Thus,

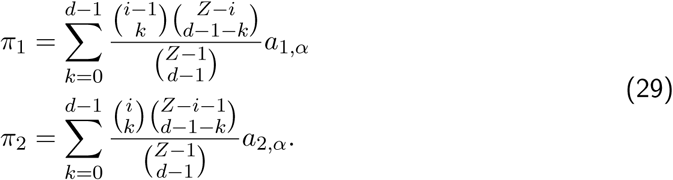

Maintaining weak selection, then from [Gokhale and Traulsen, 2010] we have,

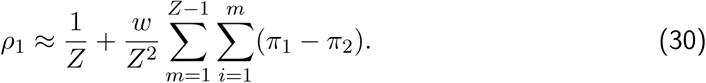

#### 2.2 Multiple game dynamics

We begin with the same example that was used to explain the combination of two 2-player games and use the same notations for a finite population of size *Z*. The population consists of individuals of four types :
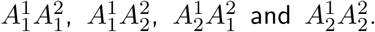 The combined dynamics results in an *S*_4_ simplex. We perform pairwise comparisons for all the edges of the simplex. On a particular edge, only the two vertex strategies are present. Let us start with the edge containing *x*_11_ and *x*_12_ vertices. If there are *γ*_11_ individuals playing strategy
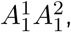 then there are *γ*_12_ = *Z* – *γ*_11_ individuals of type
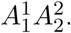 The number of
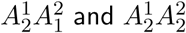 individuals i.e. *γ*_21_ and *γ*_22_ is zero. In the individual games, the number of players adopting strategy *i_j_* in a game *j* is given by *pji_j_*. Since we are looking at the edge with
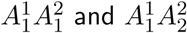 individuals, we have

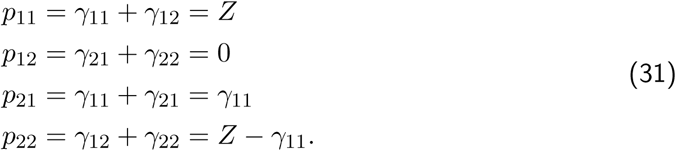

In contrast to the binomial distribution which is used for infinite populations where the draws can be considered independent, the hypergeometric distribution was used for sampling without replacement in the case of finite populations [Hauert et al., 2007, Gokhale and Traulsen, 2010]. For infinite population, we used the multinomial distribution to calculate the average payoffs for a combination of *N* multiplayer games in an infinite population size. Therefore, for finite populations, we shall use the multivariate hypergeometric distribution. For a population of size *Z* containing *γ*_11_ type
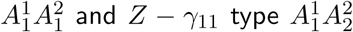 individuals, the average payoffs *π_ji_j__* for playing strategy *i_j_* in game *j* (in our example, *i_j_* ∈ {1,2} and *j* ∈ {1, 2}) are

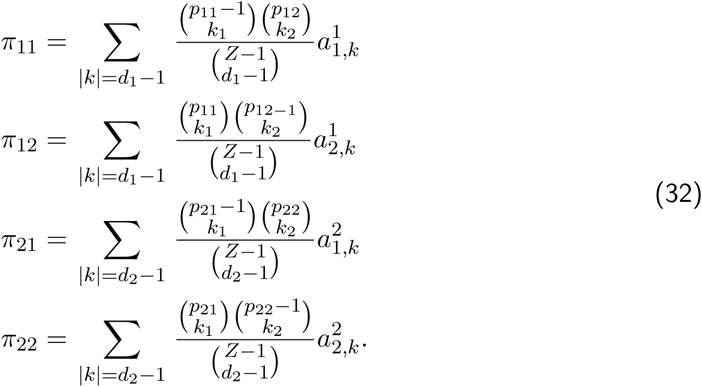

In general, for *N* multi-strategy *d*-player games,

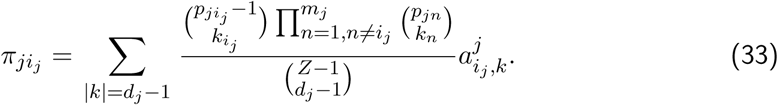

We can calculate the fitnesses using linear or exponential mapping. If *W_j_* is the intensity of selection in game *j*, then

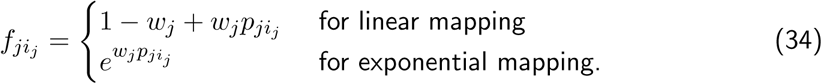

Thus, in the combined dynamics, the fitness (assuming it to be additive) of type
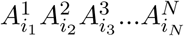
is

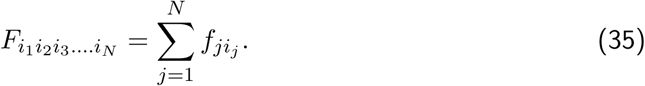

If we are looking at an edge with types
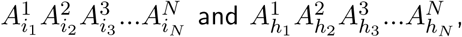
the transition probability
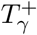
for type
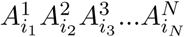
to increase from *γ* to *γ* + 1 (and type
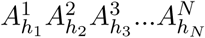
to be randomly selected for death) is

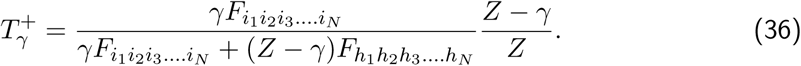

Likewise,
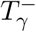
will be

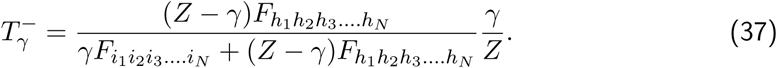

So, for a
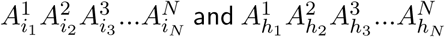
edge, the fixation probability
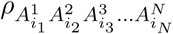
of type
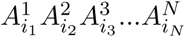
is

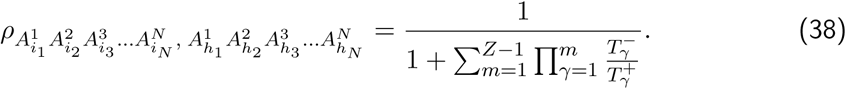

##### Result I

As
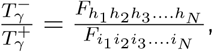
Eq. (38) can be written as

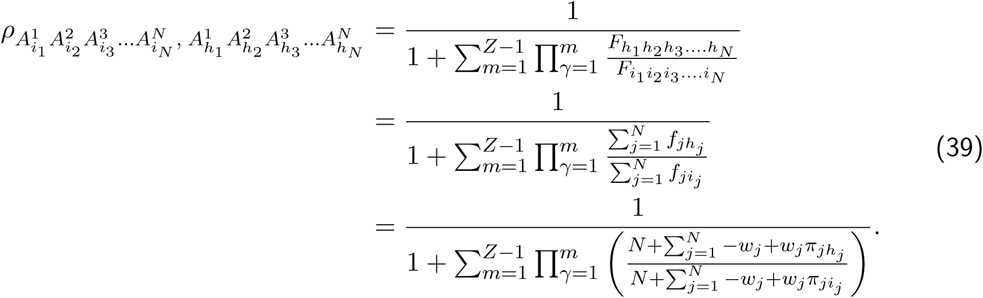

where the fitness is obtained using a linear mapping. In order to further simplify the model, we consider that all games have the same selection intensity. In this case,

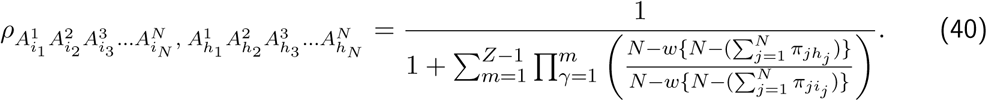

It is worth mentioning here that the assumption of having equal intensities for all games is strong. Many times, the selection on one game may be more intense than others. These have to be taken into account as it strengthens the precision of the model and Eq. (39) must be used in these scenarios. However for the sake of our analysis, we shall assume *W_j_* = *w* for all *j* ∈ [0, *N*].

For weak selection intensity,

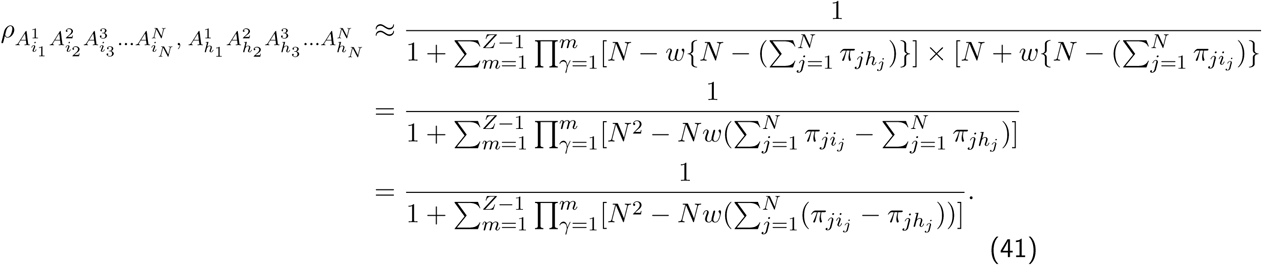

Eq. (41) can be written as

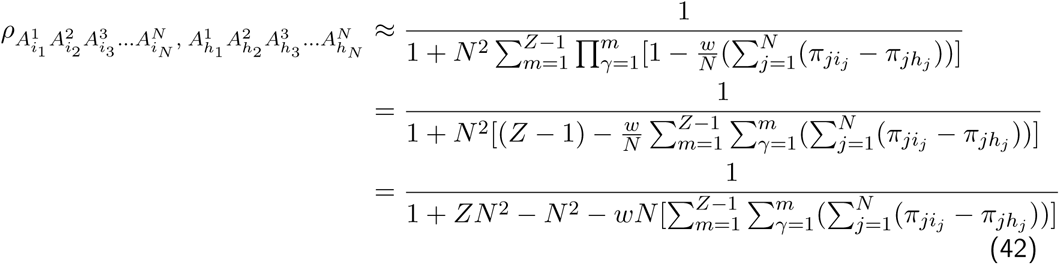

Following Taylor expansion and since *w* ≪ 1, we get

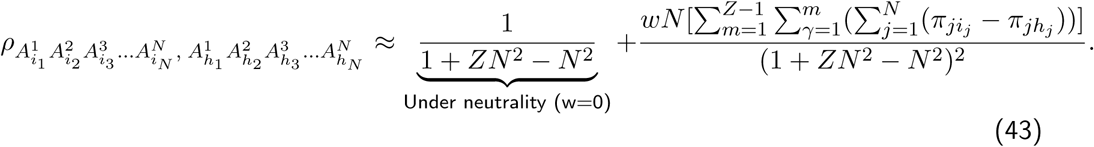

For *N* = 2 Eq. (41) becomes,

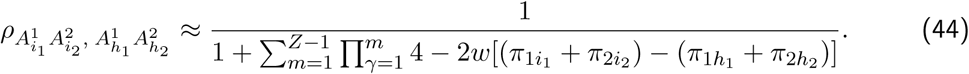

While looking at an edge for which, say, game 2 in both vertices has the same strategy and thus, we need to only look at differences in one game i.e. only game 1 matters (*π*_2*i*_2__ = *π*_2*h*_2__),

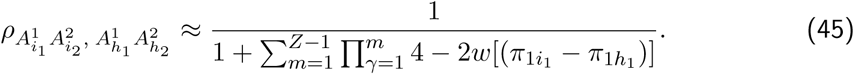

For *N* = 1 in Eq. (43), we can retrieve Eq. (30) for a single multiplayer game i.e.

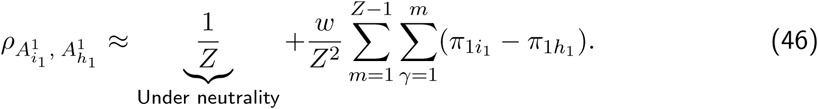

##### Result II

If all games have the same intensity, we could also add the payoffs first and then perform the fitness mappings, then
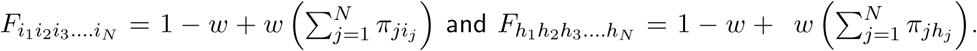
Thus, the combined fitness (of a vertex) is not just a sum of the fitnesses of strategies used in the inherent games (in that vertex). The combined fitness is obtained by summing the average payoffs of playing the respective strategies in the games involved in a particular vertex and using that to calculate the fitness of that vertex. This combination of games is not trivial as bringing all the smaller games into one larger game but we cannot always disintegrate the multi-game back to all the inherent single games.

The fixation probability Eq. (38), in this case will be,

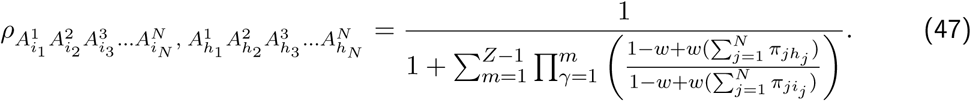

For weak selection intensities,

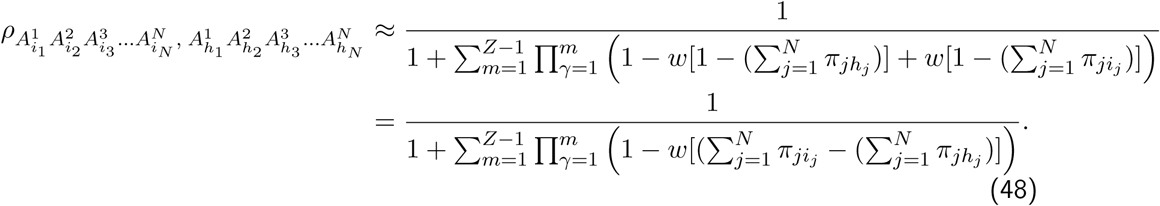

If we consider two games, then Eq. (48) will be reduced to

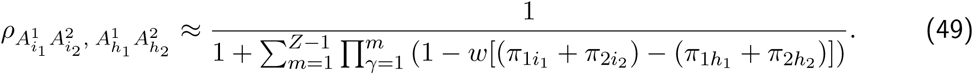

Here, if we look at an edge for which, say, game 2 in both vertices has the same strategy (*π*_2*i*_2__ = *π*_2*h*_2__), then looking at differences in game 1 is what matters. In this scenario,

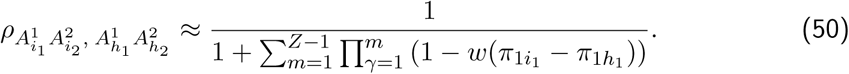

This corresponds to equation Eq. (27) for a single game with two strategies *i*_1_ and *h*_1_.

## Acknowledgments

The authors thank Peter Czuppon and Christoph Hauert for helpful discussions. Generous funding from the Max Planck Society is gratefully acknowledged.

